# Type II Alexander disease caused by splicing errors and aberrant overexpression of an uncharacterized GFAP isoform

**DOI:** 10.1101/842229

**Authors:** Guy Helman, Asako Takanohashi, Tracy L. Hagemann, Ming D. Perng, Marzena Walkiewicz, Sarah Woidill, Sunetra Sase, Zachary Cross, Yangzhu Du, Ling Zhao, Amy Waldman, Bret C. Haake, Ali Fatemi, Michael Brenner, Omar Sherbini, Albee Messing, Adeline Vanderver, Cas Simons

**Author notes:** equal contributions (Guy Helman, Asako Takanohashi). shared senior authorship (Albee Messing, Adeline Vanderver, Cas Simons). corresponding authors: Adeline Vanderver;, Cas Simons.

## Abstract

Alexander disease results from gain of function mutations in the gene encoding glial fibrillary acidic protein (GFAP), an intermediate filament protein expressed in astrocytes. At least eight *GFAP* isoforms have been described, however, the predominant alpha isoform accounts for approximately 90% of GFAP protein in the central nervous system. Here we describe exonic variants identified in three unrelated families with Type II Alexander disease that alter the splicing of *GFAP* pre-mRNA and result in upregulation of a previously uncharacterized GFAP lambda isoform (NM 001363846.1). Affected members of Family 1 and Family 2 shared the same missense variant, NM 001363846.1:c.1289G>A;p.(Arg430His) while in Family 3 we identified a synonymous variant in the adjacent nucleotide, NM 001363846.1:c.1290C>A;p.(Arg430Arg). Using RNA and protein analysis of brain autopsy samples, and a mini-gene splicing reporter assay, we demonstrate both variants result in upregulation of the lambda isoform. We assessed other *GFAP* variants in the ClinVar database for predicted aberrant splicing and using the same assay demonstrated significant changes to splicing for two selected variants. Our approach demonstrates the importance of characterizing the effect of *GFAP* variants on mRNA splicing in order to inform future pathophysiologic and therapeutic study for Alexander disease.

## Main Text

Glial fibrillary acidic protein (GFAP), encoded by *GFAP* (OMIM: 137780), is an intermediate filament protein that is highly expressed by astrocytes in the central nervous system. Pathogenic variants in *GFAP* are associated with Alexander Disease (AxD [OMIM:203450]), an autosomal dominant leukodystrophy characterized primarily by astrocyte dysfunction [17]. The majority of AxD associated variants are missense variants in coding regions of the predominant GFAP isoform, GFAP-alpha (*GFAP-α*) and are thought to exert a dominant toxic gain of function. To our knowledge, there have only been three reported cases of AxD resulting from variants affecting *GFAP* mRNA splicing [3–5].

AxD is divided into two forms based on the age of onset and symptom complex [15,18]. AxD Type I is an early childhood onset leukodystrophy with macrocephaly, spasticity, seizures, psychomotor regression, and bulbar dysfunction eventually culminating in early childhood demise. Type II AxD is typically seen in older children and adults, with a clinical presentation of autonomic dysfunction, gait dysfunction, and bulbar symptoms including palatal tremor/myoclonus and oculomotor abnormalities. Type II AxD has a more favorable prognosis, with a median survival of 25 years versus 14 years for type I AxD [15].

In this study, we report three families with a clinical presentation consistent with Type II AxD that have variants in a shared exon of the epsilon isoform of *GFAP* and a previously uncharacterized *GFAP* isoform (NM 001363846.1). Family 1 and Family 2 share the same missense variant, NM 001363846.1:c.1289G>A; p.(Arg430His), while Family 3 has a synonymous variant in the adjacent nucleotide, NM 001363846.1:c.1290C>A; p.(Arg430Arg). Using a mini-gene splicing reporter assay together with RNA and protein analysis of brain autopsy samples, we demonstrate that both variants result in aberrant splicing of *GFAP* pre-mRNA.

Informed consent for all participating individuals was obtained under IRB approvals by the Institutional Review Board at the Children’s Hospital of Philadelphia. Protein expression studies were performed under institutional review board approvals from the University of Alabama at Birmingham, Albert Einstein College of Medicine, and the University of Wisconsin-Madison.

Family 1 has a five-generation pedigree that includes more than 20 affected individuals with a phenotype consistent with Type II AxD. Analysis of their clinical data (Table 1) suggests that affected individuals presented most commonly with gait dysfunction (12/16), bulbar dysfunction manifesting as swallowing difficulties, choking, or dysarthria (7/16), and extremity paresthesias (4/16). The average age of onset was 42 years of age (range: 20 – 66 years). MRI, performed in 6 individuals, showed T_2_ hyperintensity in the brainstem in all cases, often affecting the medulla (6/6). Significant atrophy of the medulla was seen in two of these. Autopsy performed in two affected individuals demonstrated the hallmark neuropathologic features of AxD, including Rosenthal fibers and astrogliosis. A constellation of clinical tests indicative of a cause of their neurologic disease across different members of this family was unrevealing, yet the family history of dominantly-inherited disease and clinical features including neuropathologic confirmation consistently suggested a diagnosis of AxD (Figure 1A and Table 1).

**Table 1.**
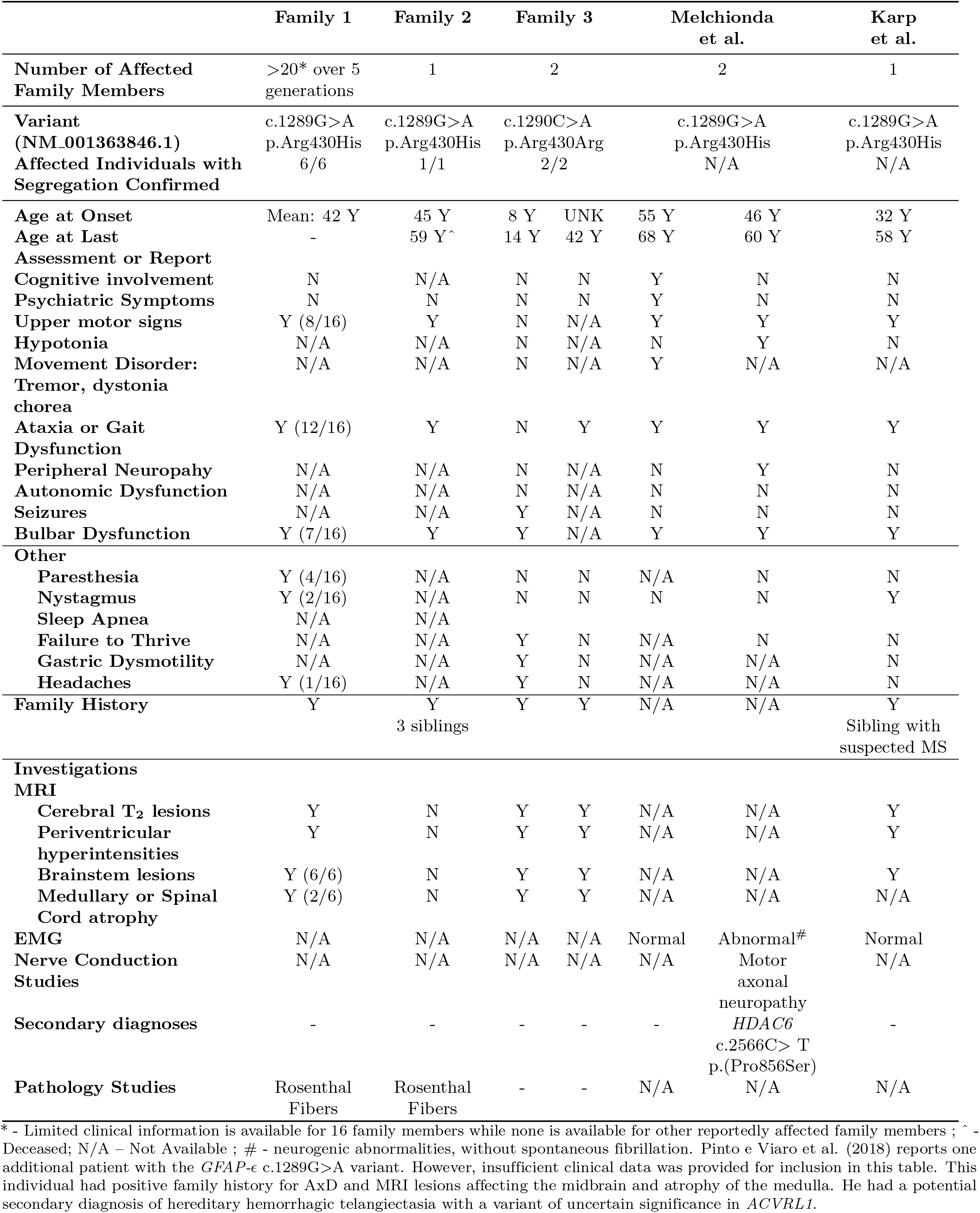
Clinical characteristics

**Figure 1.**
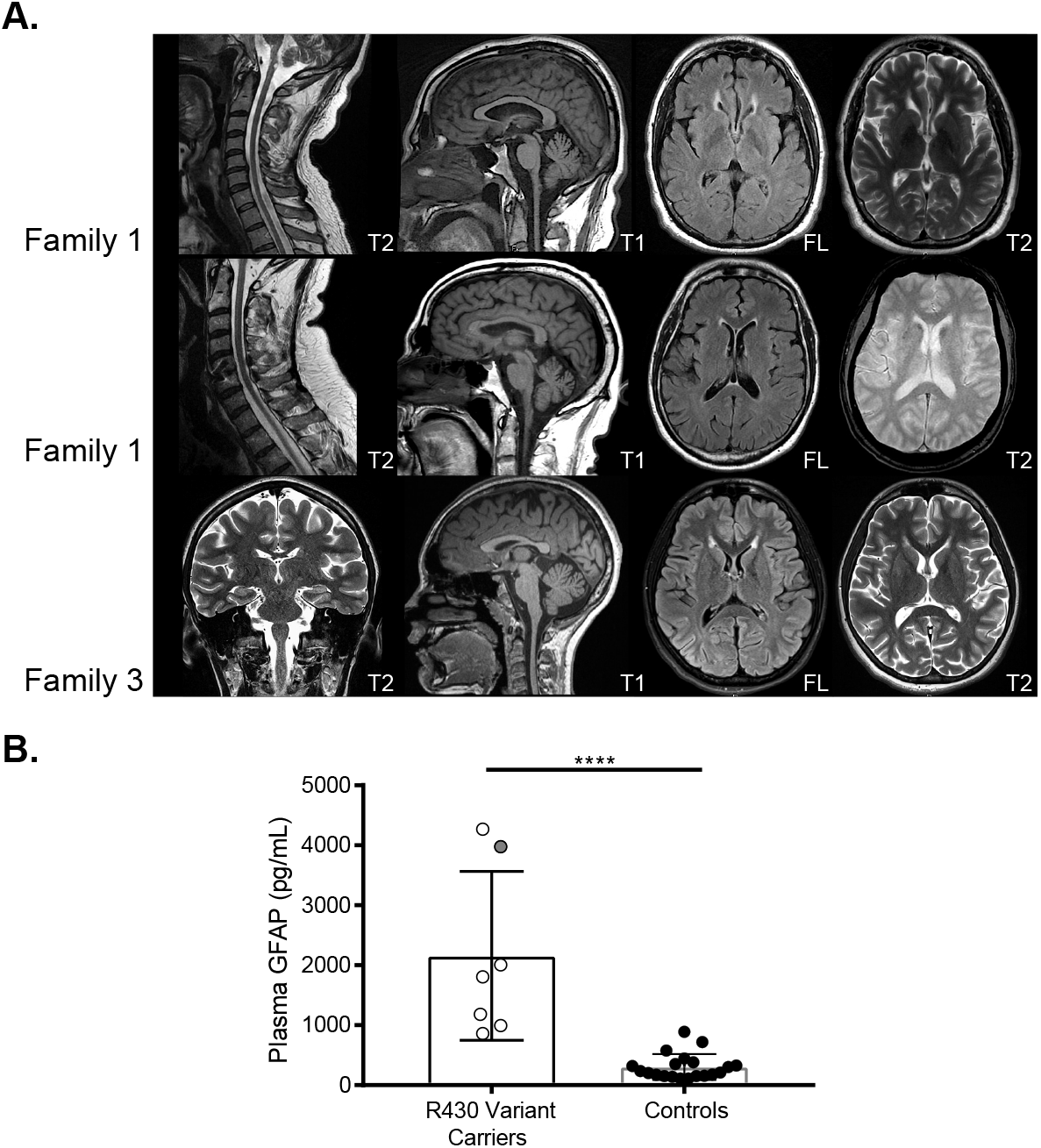
MRI and plasma GFAP levels of select affected individuals. (A) MRI of selected affected individuals. Imaging of two separate affected individuals from the same nuclear family of Family 1 (Row 1 aged 48 years and Row 2 aged 62 years) shows medullary atrophy. In all affected individuals there are periventricular hyperintensities in the region of the frontal horns of the lateral ventricles on FLAIR and T_2_ sequences. Imaging from Individual 1 in Family 3 (Row 3) was performed at 13 years of age. (B) Plasma GFAP levels were determined by digital ELISA. As a group, six individuals heterozygous for either the c.1289G>A (annotated in white with black outline) and a single individual with the c.1290C>A (annotated in grey with black outline) variants demonstrated significantly elevated blood plasma GFAP levels relative to unaffected controls (p=0.0001).

Family 2 consists of an affected male previously reported as Patient 43 in Li et al. (2005) [10], with a clinical presentation suggestive of adult-onset AxD, that was confirmed by autopsy, but for whom a causative *GFAP* variant had not been detected. He presented at 45 years of age with gait dysfunction, dysarthria, and upper motor neurological signs. There was also significant family history, with three siblings found to also be clinically affected with features consistent with Type II AxD.

Family 3 consists of a female proband and her affected father, with a disease phenotype suggestive of Type II AxD (Table 1). The proband had disease onset at 10 years of age, with headaches, bulbar dysfunction, failure to thrive, gastric dysmotility, and seizures. MRI was consistent with AxD (Figure 1A), with notable T_2_ signal abnormalities in the brainstem (medulla and pons) and the frontal horns of the lateral ventricles as well as mass-like lesions along the brainstem. The father’s MRI demonstrated a brainstem lesion and he has had increasing difficulties in gait and swallowing, as well as autonomic dysfunction, dysarthria and sleep apnea as commonly seen in Type II AxD.

Targeted sequencing of *GFAP* was performed on one affected individual from Family 1 and the probands from Families 2 and 3. The tested individuals from Family 1 and Family 2 were both found to be heterozygous for the same missense variant NM 001363846.1:c.1289G>A;p.(Arg430His). The proband from Family 3 was heterozygous for a synonymous variant NM 001363846.1:c.1290C>A; p.(Arg430Arg) in the nucleotide adjacent to the c.1289G>A variant in identified in Families 1 and 2.

Segregation analysis by Sanger sequencing confirmed that all six tested members of Family 1 with clinical symptoms were heterozygous for the c.1289G>A variant. In Family 3, segregation analysis confirmed the proband’s affected father was also heterozygous for the c.1290C>A variant.

In healthy central nervous system tissue, approximately 90% of GFAP protein is derived from the predominant GFAP-*α* isoform (NM 002055.5) [12]. At least seven other *GFAP* isoforms have also been described, including the GFAP epsilon isoform (GFAP-*ϵ*; NM 001131019.3), also known as the delta isoform (GFAP-*δ*); as well as the kappa isoform (GFAP-*κ*; NM 001242376.2) [1]. The initial 390 amino acids are identical in all major GFAP isoforms, and differ only in their make-up of the C-terminal region (Figure 2A-B).

**Figure 2.**
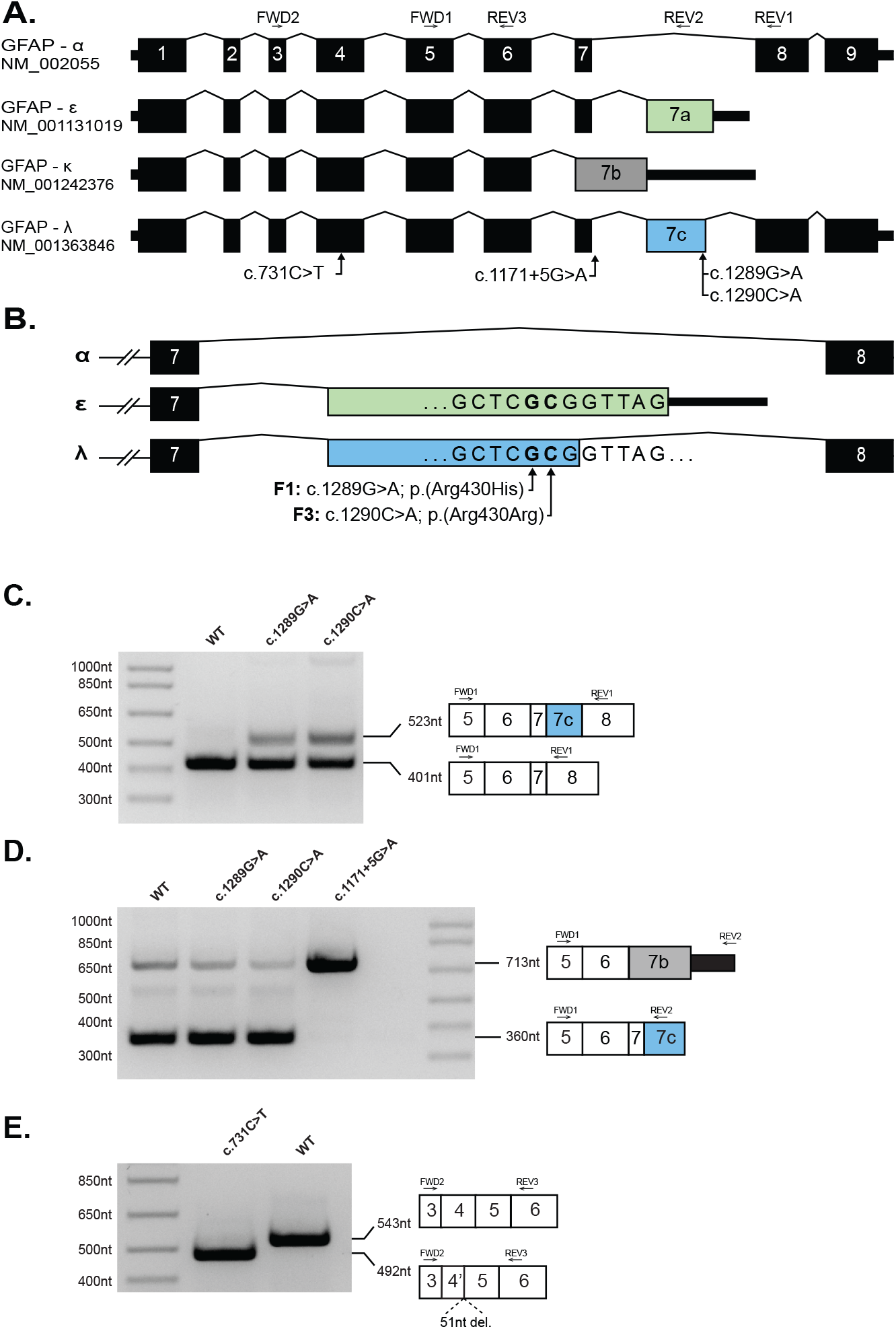
Missense and synonymous variants in GFAP result is alteration in GFAP pre-mRNA splicing. (A) Schematic of four *GFAP* pre-mRNA isoforms. Exons 1-9 are numbered per the *GFAP-α* isoform. Introns are not shown to scale. (B) Schematic of C-terminal variants between exons 7-8 of *GFAP* showing the positions of the single nucleotide substitutions identified in Family 1 (F1) and Family 3 (F3) relative to exon 7A (green, *GFAP-ϵ*) and exon 7C (blue, *GFAP*-λ). (C) A GFAP mini-gene plasmid was created by introducing 8.1kb of *GFAP* genomic DNA (exon1 – exon 9, amino acids 57-432) into the mammalian expression vector pEYFP-C1 (see Supplemental Methods). Versions of this construct were generated containing the *GFAP-ϵ* variants, c.1289G>A and c.1290C>A, or the *GFAP-α* variants c.731C>T and c.1171+5G>A variants before transfection into HEK cells. rtPCR amplification from exon 5 to exon 8 yielded a predominant amplicon consistent with the *GFAP-α* isoform (401nt) from the WT (Lane 1) c.1289G>A (Lane 2) and c.1290C>A (Lane 3) constructs. The c.1289G>A and c.1290C>A mutant constructs also generated a larger amplicon consistent with the *GFAP*-λ isoform (523nt). (D) Amplification from exon 5 to exon 7A/7C yielded a predominant amplicon consistent with the *GFAP-ϵ* and *GFAP*-λ isoforms (360nt) from the WT (Lane 1), c.1289G>A (Lane 2) and c.1290C>A (Lane 3) constructs, and a minor amplicon consistent with the *GFAP-κ* (713nt) is also visible in each of these lanes. The c.1171+5G>A mutant construct yielded a single predominant amplicon consistent in size with *GFAP-κ* (713nt). (E) Amplification from exon 3 to exon 5 (common to all major *GFAP* isoforms) yielded a single amplicon consistent with the expected size of 543nt from the WT construct (Lane 2). Amplification from the construct containing the c.731C>T missense variant (Lane 1) yielded a smaller single amplicon consistent with a truncation of exon 4. Sanger sequencing of the c.731C>T amplicon confirmed a 51nt truncation of exon 4 (Supplemental Figure S2).

The c.1289G>A and c.1290C>A variants identified in this manuscript are both situated in an exon shared by *GFAP-ϵ* (exon 7A) and the transcript NM 001363846.1, hereafter referred to as *GFAP*-λ (Figure 2A-B). The *GFAP*-λ transcript is present in the RefSeq mRNA database [14], but to our knowledge has not previously been described in the AxD or GFAP literature. The exon containing c.1289G>A and c.1290C>A is not included in the predominant *GFAP-α* isoform.

The c.1289G>A variant has been previously described in several cases; a sibling pair with adult-onset AxD [11], and two descriptions of single, unrelated individuals in Karp et al. and Pinto e Vairo et al. [9, 13] (Table 1). It is possible that the patient reported by Pinto e Vairo et al. (2018) is related to the individuals we have studied in Family 1, and hence not an independent occurrence of this variant [13]. In the study by Melchionda et al., it was hypothesized that the pathogenic effect of the c.1289G>A variant resulted from perturbation of the GFAP network by inefficiently incorporated GFAP-*ϵ*^Arg430His^ protein [11].

Single Molecule Array (Simoa; Quanterix), a digital ELISA assay, was used to quantitate GFAP levels in blood plasma of individuals from both Family 1 and Family 3 Individuals affected by AxD are known to have highly elevated levels of GFAP in plasma [8]. We found individuals heterozygous for either the c.1289G>A (Family 1) or the c.1290C>A (one individual in Family 3) variants demonstrated significantly elevated blood plasma GFAP levels relative to controls (p=0.0001) (Figure 1B).

Given that Families 1 and 2 have a single nucleotide variant adjacent to that of Family 3, only the former of which is predicted to result in a change to an encoded protein sequence, we hypothesized that these variants may have a deleterious impact through influencing the splicing of *GFAP* pre-mRNA. To address this possibility, we generated a mini-gene splicing reporter by cloning 8.1kb of GFAP genomic sequence into the mammalian expression vector pEYFP-C1. Primer directed mutagenesis was used to generate versions of the plasmid containing the c.1289G>A and c.1290C>A variants before transfection into HEK cells.

Amplification by reverse transcription-PCR (RT-PCR) using a forward primer in exon 5 and a reverse primer in exon 8 yielded an amplicon consistent with the size expected from *GFAP-α* using RNA isolated from cells transfected with either the wild-type or the mutant plasmids (Figure 2C). However, an additional, approximately 100 nt larger amplicon was also generated from the cells transfected with either the c.1289G>A or c.1290C>A mutant plasmids that is absent from cells transfected with the wild-type plasmid (Figure 2C). Sanger sequencing revealed that the larger amplicon is due to the presence an additional exon, hereafter referred to as exon 7C, between exon 7 and exon 8, consistent with the *GFAP*-λ isoform (Figure 2C and Supplemental Figure S1).

RNA extracted from brain autopsy samples from a deceased member of Family 1 was used to confirm the presence of the aberrantly spliced transcript. Primers located in exons 6 and 9 of *GFAP-α* amplified a fragment of the expected size for a *GFAP-α* template from both patient and control brain RNA samples (Figure 3A). The patient RNA also generated a larger amplicon consistent with the *GFAP*-λ splicing variant noted above (Figure 3A). Using a primer pair specific to *GFAP*-λ, we confirmed its presence in the patient sample. The *GFAP*-λ splice variant was also detected in the control sample, although it was noticeably less abundant (Figure 3A). Sanger sequencing confirmed the identity of these amplicons (Figure 3B).

**Figure 3.**
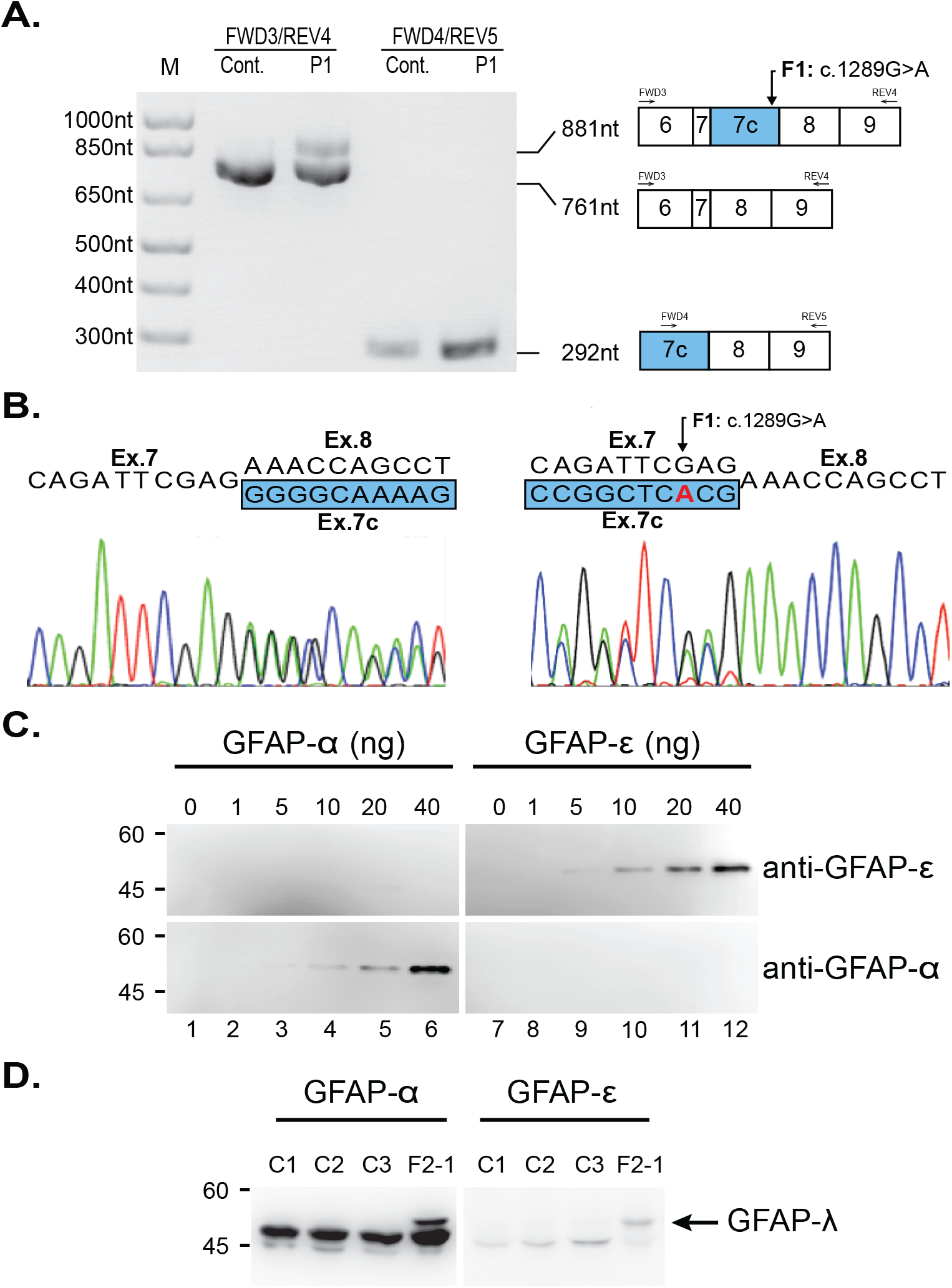
Demonstration of the presence of *GFAP*-λ transcript and protein in the AxD brain. (A) RT-PCR of mRNA from autopsy brain tissue of an affected member of Family 1 (F1-1) and an unaffected control. Lanes 1 and 2: amplification between exons 6 and 9 generates an amplicon of the expected 761nt in both the patient and control cDNA samples. An additional 881nt amplicon can be seen in the cDNA sample of the affected individual (F1-1), consistent with *GFAP*-λ. Lanes 3 and 4: amplification between exons 7C and 9 generates a *GFAP*-λ specific amplicon of 292nt that is detectible in the control cDNA sample but noticeably more abundant when amplified from the patient cDNA. (B) Sanger sequencing of the mixed amplicon from lane 2 in both forward (left) and reverse (right; reverse complement shown) shows the expected nucleotide sequences and splice sites for *GFAP-α* and *GFAP*-λ. (C) The specificity and affinity of the isoform-specific GFAP antibodies were characterized by immunoblotting using dilutions of known concentrations of purified recombinant human GFAP-*α* (lanes 1-6) and GFAP-*ϵ* (lanes 7-12). The anti-GFAP-*α* and anti-GFAP-*ϵ* antibodies recognized purified GFAP-*α* (lanes 4-6, bottom panel) and GFAP-*ϵ* (lanes 9-12, top panel), respectively, confirming the specificity of these antibodies. (D) Intermediate filament-enriched fractions were prepared from brains of four AxD affected individuals (lanes 1-4), including an affected family member of Family 2 (F2-1, lane 4). Samples (10-12 *μ*g protein load per lane) were analyzed by immunoblotting with anti-GFAP-*α* (lanes 1-4) and anti-GFAP-*ϵ* (lanes 5-8) antibodies. Note the presence of the GFAP-λ in F2-1 (lanes 4 and 8, arrow).

The *GFAP*-λ transcript is equivalent to *GFAP-α* with an additional 120nt exon (exon 7C) inserted between exons 7 and 8 (Figure 2A). Exon 7C, shares the same splice acceptor site as the terminal exon of *GFAP-ϵ* (exon 7A), but utilizes a splice donor site 2 nt 5’ of the stop codon of *GFAP-ϵ*. The predicted protein encoded by *GFAP*-λ consists of the first 430 (of 431) residues of GFAP-*ϵ* followed by the terminal 42 residues GFAP-*α*.

To determine if the *GFAP*-λ mRNA is translated into a stable protein, we performed western blot analysis on protein isolated from a brain autopsy sample from the proband of Family 2. The blot was probed with an available antibody to an epitope in exon 9 of GFAP-*α*, which should recognize GFAP-*α* and GFAP-λ, but not GFAP-*ϵ*, and a separate antibody to an epitope in exon 7A of GFAP-*ϵ*, which should recognize GFAP-*ϵ* and GFAP-λ, but not GFAP-*α*. Blots confirming the specificity of these antibodies were performed with recombinant GFAP-*α* and GFAP-*ϵ* (Figure 3C). In intermediate filament enriched protein fractions, the expected 50 kD GFAP-*α* and 49.5 kD GFAP-*ϵ* are clearly present in the Family 2 patient as well as three unrelated AxD controls (Figure 3D). In the individual from Family 2, a larger band is clearly visible that appears to cross react with both antibodies. This band is not present in any of the other AxD affected samples serving as controls and is consistent with the predicted protein encoded by GFAP-λ (54 kD) that contains the C-terminal regions of both GFAP-*α* and GFAP-*ϵ*.

Based on the hypothesis that alteration of splicing might be a broader mechanism of pathogenesis in AxD, we reviewed *GFAP* variants recorded in ClinVar and used SpliceAI [7] to predict if they were likely to alter splicing, identifying two variants strongly predicted to do so. NM 002055.5:c.1171+5G>A (ClinVar ID:323610; located 5 nt into the start of *GFAP-α* intron 7), is predicted to weaken the exon 7 splice donor site, which could favor formation of the kappa isoform (Figure 2B). The second variant, NM 002055.5:c.731C>T;p.(Ala244Val) (ClinVar ID: 66500), has been associated with Type I AxD and classified as pathogenic on the basis of clinical findings despite inconclusive functional testing [10]. SpliceAI predicts this variant will strengthen a cryptic splice donor site within exon 4, truncating the final 17 amino acids of the exon while maintaining the reading frame. We interrogated each of these variants using our GFAP mini-gene splicing reporter assay and confirmed that each of the variants induced the predicted change in *GFAP* splicing (Figure 2C-E and Supplemental Figure S2).

In this study, we demonstrate that the *GFAP* variants c.1289G>A and c.1290C>A both result in a substantial alteration in the splicing of *GFAP* and an upregulation of a previously undescribed isoform, GFAP-λ. Previous *in vitro* studies have shown that overexpression of GFAP-*ϵ* induces changes to intermediate filament networks and increases associations with cellular stress proteins such as *α*B-crystallin [12]. In Vanishing White Matter Disease (OMIM:603896), an autosomal recessive leukodystrophy also characterized by astrocytic dysfunction, *in vivo* models and neuropathology have revealed significantly increased GFAP-*ϵ* and dysmorphic astrocytes, likely affecting trophic support to developing oligodendrocytes [2, 19]. Given GFAP-*ϵ* is a truncated version of GFAP-λ, we suggest that the increase in GFAP-λ is likely to result in a similar perturbation of the GFAP network. Although we cannot discount the possibility that the p.(Arg430His) missense change in GFAP-*ϵ* is also deleterious (as suggested by Melchionda et al. (2013)), our finding of an adjacent synonymous c.1290C>A; p.(Arg430Arg) variant in Family 3 suggests that alteration of *GFAP* splicing is likely to be the primary functional consequence of both of these AxD associated variants.

Type II AxD is characterised typically by a later age of onset but can be variable, as evidenced by the juvenile onset disease seen in an affected individual of Family 3. Thus, evaluating the penetrance of particular pathogenic variants can be challenging. A full understanding of the penetrance of the variants discussed in this manuscript will require larger segregation studies and longer longitudinal follow-up.

It was recently demonstrated that antisense suppression using targeted oligonucleotides in a murine model of AxD attenuated and reversed the overexpression of *Gfap* and lowered tissue and cerebrospinal fluid GFAP protein levels, providing a promising therapeutic approach for this disorder [6]. Antisense approaches have demonstrated benefit in other disorders with a gain-of-function mechanism, and appropriate design must take into account variant consequences, including aberrant splicing [16]. The results presented in this manuscript suggest that selective suppression of individual isoforms might be a future avenue for therapeutic development for AxD cases associated with aberrant splicing.

In summary, while the majority of variants causing AxD are missense variants in coding regions, we provide further evidence that aberrant splicing due to exonic and intronic variants in *GFAP* is likely to contribute to a fraction of AxD cases. Our study illustrates the importance of considering the effect of *GFAP* variants on mRNA splicing to ensure accurate diagnosis, and to appropriately plan future therapeutic approaches for affected individuals.

## Acknowledgments

The authors would like to thank the affected individuals and their families. We would like to thank Professor Elly M. Hol for contributing antibodies used in this study, Johanna Schmidt for her work with the Myelin Disorders Bioregistry Project, and would like to acknowledge the Human Immunology Core (P30-CA016520) at the University of Pennsylvania for the Simoa assays performed in this study.

## Supporting Information

A supplemental text file has been provided and includes a full description of methods.

