## Supplemental Text for "Type II Alexander disease caused by splicing errors and aberrant overexpression of an uncharacterized GFAP isoform"

### To whom correspondence should be addressed
* shared first authorship
** shared senior authorship

**Supplemental Methods:**

*Patient Selection*

Informed consent for all participating individuals was obtained under IRB approval by the Institutional Review Board at the Children's Hospital of Philadelphia (IRB#14-011236). Protein expression studies were performed under institutional review board approvals from the University of Alabama at Birmingham, Albert Einstein College of Medicine, and the University of Wisconsin-Madison. Available patient medical records and neuroimaging were reviewed.

*Plasma GFAP Screening*

GFAP protein levels were tested from plasma samples isolated from patients, unaffected family members, and controls using an ultrasensitive Single Molecule Array (Simoa, also called digital ELISA, Quanterix™, #102336) GFAP immunoassay as previously described (Abdelhak, et al., 2018). A one-way ANOVA with multiple comparisons was used for multigroup comparisons and for two group comparisons an unpaired t-test was performed and plotted using Prism software (Graphpad, Inc.).

*Clinical Sequencing and Segregation Analysis*

Single gene sequencing was performed in separate CLIA-approved laboratories for each family. Segregation analysis for Family 1 was performed by Sanger sequencing of available blood DNA samples on an Applied Biosystems 3730XL system (Thermo Fisher Scientific).

*Mini-gene splicing assay*

Mini-gene splicing assays were employed to assess the splicing impact of four *GFAP* variants predicted to impact splicing. We cloned the majority of genomic sequence of *GFAP* (Exons 1-9, Amino Acids 57-433) and introduced variants using three fragment HiFi DNA Assembly. The mutant-bearing fragments were isolated using the Monarch DNA Gel Extraction Kit (New England BioLabs, #T1020L) and introduced into a linearized EYFP-C1 vector using the NEBuilder HiFi DNA Assembly Kit (New England BioLabs, #E5520S) (see Table S1 for primers), resulting in an insert size of approximately 8.1 kb and a total vector size of 12kb.

The assembled product was transformed using NEB 5-alpha Competent *E. coli* cells and plated on an LB agar plate containing kanamycin. Plasmids for each mutant and wild-type construct were Sanger-sequenced to confirm the presence of each variant in mutated constructs and the absence of variants in the wild-type construct (see Table S2 for primers).

Wild-type and mutant plasmids were transfected into the HEK293T cell line (Lipofectamine™ 2000 reagent (Thermo Fisher Scientific, #11668019)) and cultured for 48 hours. RNA was isolated from the transfected cells using the RNeasy Plus Mini Kit (Qiagen) and cDNA was synthesized using a High-Capacity cDNA Transcription Kit (Applied Biosystems) according to the manufacturer's instructions. RT-PCR was performed using *GFAP* exonic primers (see Table S1 for primers). The resulting bands were purified using the Monarch DNA Gel Extraction Kit and cloned using the TOPO TA Cloning kit (Thermo Fisher Scientific, #K4500-01 before Sanger sequencing (see Table S2 for primers).

*RT-PCR from Patient Brain*

RNA was isolated from the brain tissue sample of a deceased member of Family 1 carrying the c.1289G>A variant. RNA was isolated using the RNeasy Lipid Tissue Mini Kit (Qiagen # 74804) and treated with DNase I. cDNA was then generated using the SuperScript III First-Strand Synthesis System (Invitrogen #18080-051). RT-PCR was performed using Q5 Hot Start High-Fidelity DNA polymerase (New England BioLabs #M0493) and gene-specific primers (Table S3) for 35 cycles. The resulting PCR products were run on E-Gel Size Select II Agarose Gel, 2% (Life Technologies # G661012), isolated and purified using Genomic DNA Clean & ConcentratorTM-10 (Zymo Research, #D4010). The final PCR products were purified with ExoSAP-IT (Life Technologies #78201.1mL). Purified PCR products were analyzed by Sanger sequencing (see Table S2 for primers).

*Human brain samples*

Clinical and genetic details of human brain samples used for protein studies are presented in Table S4. AxD human tissues were obtained from the NIH Neurobiobank with approval from the institutional review board at the University of Wisconsin-Madison.

*Preparation of the intermediate filament-enriched fractions*

Brain tissues (0.05-0.1 mg) were thawed on ice and dounce-homogenized in low salt buffer (20 mM Tris-HCl, pH 7.4, 5 mM EDTA, 1% (v/v), Triton X-100, and 150 mM KCl). All buffers were supplemented with a cocktail of protease inhibitors, containing 10 µM ALLN, 2 µg/ml leupeptin, 5 µg/ml aprotinin, and 2 mM PMSF. Homogenates were centrifuged at 14,000 rpm for 15 minutes at 4°C, and the resulting pellets were extracted in high salt buffer (20 mM Tris-HCl, pH 7.4, 5 mM EDTA, 0.5% (v/v) Triton X-100, and 1.5 M KCl). After centrifugation at 19,000 g for 15 min at 4°C, the pellets were resuspended in urea buffer (6 M urea, 10 mM Tris-HCl, pH 7.4, and 1 mM EDTA) and further extracted at 4°C overnight. The extracts were centrifuged at 19,000 g at 4°C for 5 minutes, and the urea-soluble fractions were taken as the IF-enriched fractions. After concentration determination by BCA protein assay, protein samples were resolved by 12% (w/v) SDS-PAGE using a Tris/glycine electrophoresis system prior to analysis by immunoblotting.

*Immunoblotting*

Immunoblotting was performed using the wet electrophoretic transfer system (Biorad, Hercules, CA) according to the manufacturer's instructions. Following electrophoretic transfer, the nitrocellulose membranes (Pall Corporation, Ann Arbor, MI) were washed three times with Tris-buffered saline (TBS: 150 mM NaCl, 20 mM Tris-HCl, pH 7.4), followed by blocking in TTBS ((0.1% (v/v) Tween-20 in TBS) containing 3% (w/v) bovine serum albumin (BSA)) for 1 h at room temperature. After blocking, the membranes were incubated at 4°C overnight with mouse anti-GFAP-α (clone 52; BD Biosciences) and rabbit anti-GFAP-ε antibodies (Roelofs, et al., 2005). The specificity of these antibodies was tested by immunoblotting using recombinant human GFAP-α and GFAP-ε proteins purified as described previously (Perng, et al., 2008). After being washed with TTBS, membranes were incubated with horseradish peroxidase–conjugated goat anti-mouse or anti-rabbit secondary antibodies (Jackson ImmunoResearch Laboratories) for 2 h at room temperature. All antibodies were diluted in TTBS containing 1% (w/v) BSA. Antibody labeling was detected by enhanced chemiluminescence substrate (Western Lightning Plus, PerkinElmer; Waltham, MA) with use of a luminescent image analyzer (LAS 4000; GE Healthcare).

**Table S1: Minigene assembly primers**

| Primer | Primer Sequence (5' to 3') |
|---|---|
| EYFP-C1:GFAP_ex1_fwd | gtcagatccgctagcgctaccgatgACTCAATGCTGGCTTCAAGG |
| EYFP-C1:GFAP_ex9_rev | gtttcaggttcagggggaggtgtgGAGGGGAGCAGCTGGGGTG |
| GFAP_1289G-A_fwd | GCCGGCTCaCGGTTAGCTGCCTGCCTCTC |
| GFAP_1289G-A_rev | GCAGCTAACCGtGAGCCGGCGGCGTTCC |
| GFAP_1290C-A_fwd | GCCGGCTCGaGGTTAGCTGCCTGCCTCTC |
| GFAP_1290C-A_rev | GCAGCTAACCtCGAGCCGGCGGCGTTCC |
| GFAP_WT_fwd | GCCGGCTCGCGGTTAGCTGCCTGCCTCTC |
| GFAP_WT_rev | GCAGCTAACCGCGAGCCGGCGGCGTTCC |
| GFAP_1171+5G-A_fwd | CAGATTCGAGGTCAaTACAGCAGGGCCTCG |
| GFAP_1171+5G-A _rev | GCCCTGCTGTATtGACCTCGAATCTGC |
| GFAP_731G-A_fwd | CAGTATGAGGtAATGGCGTCCAGCAAC |
| GFAP_731G-A_rev | CTGGACGCCATTaCCTCATACTGCGTGC |

**Table S2: Sanger sequencing primers**

| Primer | Primer Sequence (5' to 3') |
|---|---|
| GFAP_ex7_fwd | GGGCAAAAGCACCAAAG |
| GFAP_ex6_fwd | GAGATCGCCACCTACAGGAAGC |
| GFAP_ex3_fwd | GATTGAGTCGCTGGAGGAGG |

**Table S3: RT-PCR of Family 1 Autopsy Tissue**

| Primer | Primer Sequence (5' to 3') |
|---|---|
| FWD1 | CCAAGCACGAAGCCAACGACTAC |
| REV1 | CTCTCCATCCCGCATCTCC |
| FWD2 | GATTGAGTCGCTGGAGGAGG |
| REV2 | CCGTCTTTGGTGCTTTTGCC |
| FWD3 | GCACGAAGCCAACGACTA |
| REV3 | CTGTAGGTGGCGATCTCGATGTC |
| FWD4 | ACGGGGAAAATCACAAGGTCA |
| REV4 | TTCTCTCCTTCCTCCTCATTCT |
| REV5 | CTATCCTGCTTCTGCTCGGG |

**Table S4. Clinical and genetic details of AxD control and Family 2 patient samples**

| Case | ID Number | Disease | Age at death | GFAP mutation | Sex |
|---|---|---|---|---|---|
| C1 | #13 in Li, et al. (2005) | Adult-onset/type II | 38 y | Glu210Lys | F |
| C2 | #34 in Li, et al. (2005) | Juvenile-onset /type II | 20 y | Leu359Val | M |
| C3 | #4 in Li, et al. (2005) | Infantile-onset/type I | 10.5 y | Leu76Val | F |
| F2-1 | #43 in Li, et al. (2005) | Adult-onset/type II | 59 y | Arg430His/Arg430Cys | M |

**Supplemental Results:**

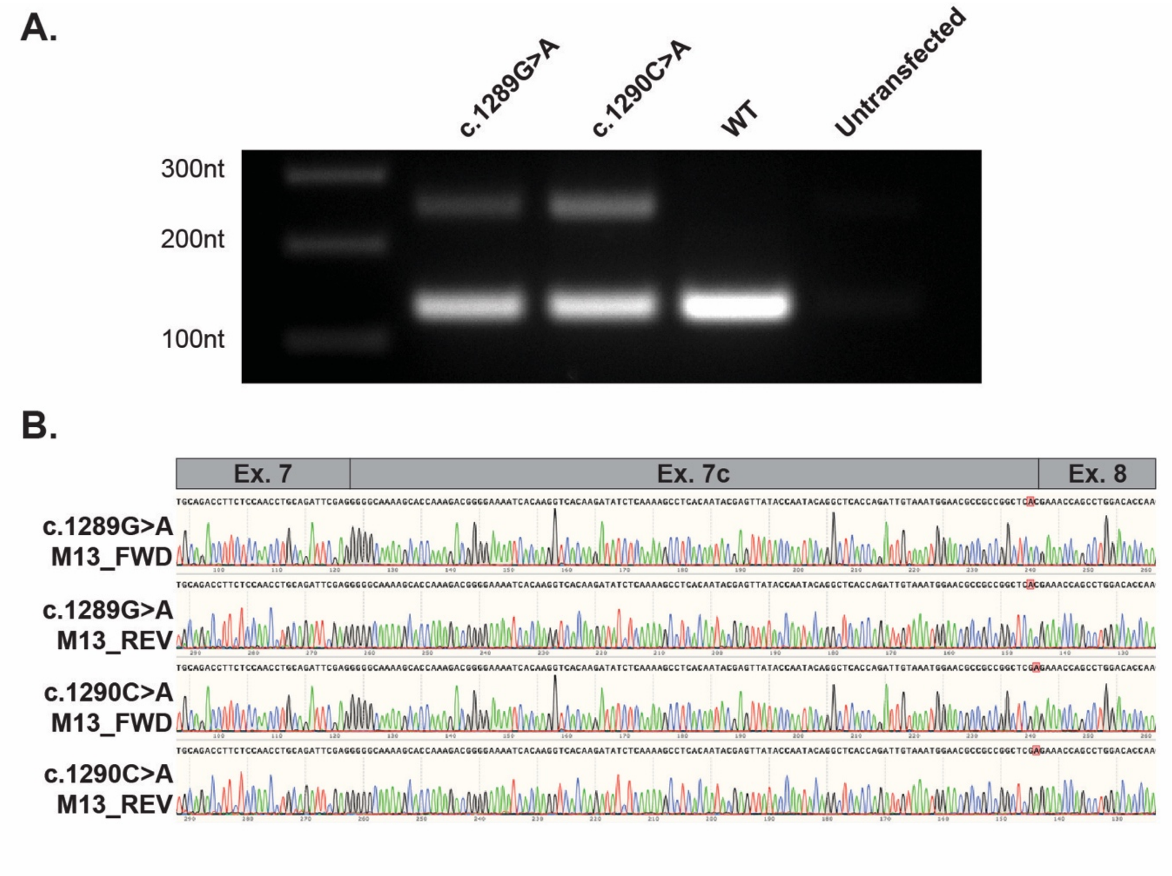


**Figure S1** – **Aberrant *GFAP* splicing due to the c.1289G>A and c.1290C>A variants results in production of the GFAP-λ isoform**. (A) RT-PCR from RNA extracted from HEK cells transfected with GFAP minigene plasmids. An amplicon consistent with the expected size for GFAP-*α* (130 nt) was generated from the c.1289G>A (Lane 1), c.1290C>A (Lane 2) and WT plasmids (Lane 3). A larger band consistent with the expected size for GFAP-λ (252 nt) is only visible from the mutant plasmids c.1289G>A and c.1290C>A. (B) Sanger sequencing of the cloned 252 nt amplicon from each mutant plasmid demonstrates inclusion of exon 7c between exon 7 and exon 8, representative of the GFAP-λ transcript. The position of each variant is highlighted in red.

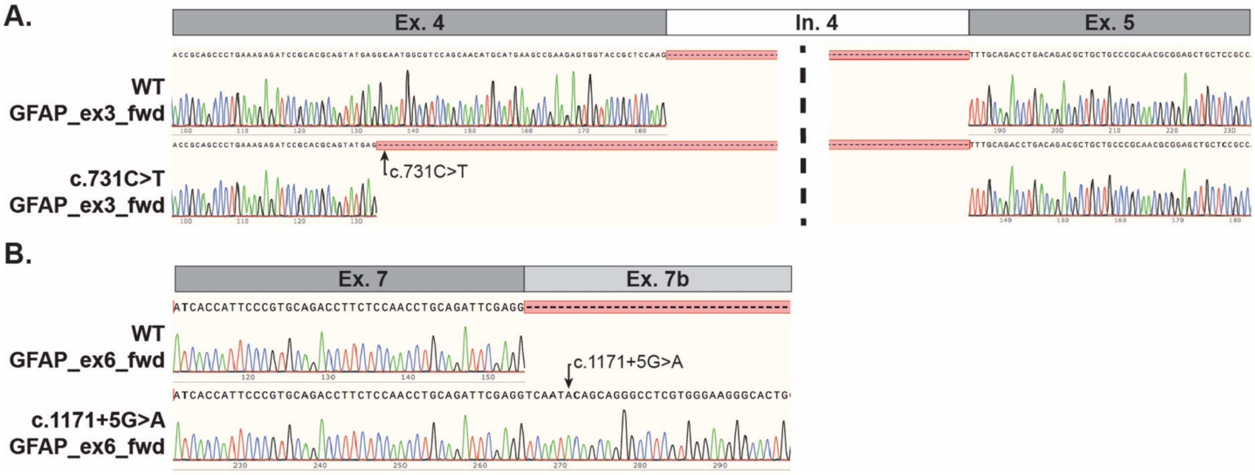


**Figure S2 -Aberrant *GFAP* splicing of mini-gene constructs bearing either the c.731C>T or the c.1171+5G>A variant.** (A) Amplicons shown in Figure 2E were gel extracted and Sanger sequenced. The c.731C>T variant results in a 51 nt truncation of Exon 4. The intervening sequence from Intron 4 is shown truncated at the dashed line. (B) The amplicon 360nt amplicon (WT) and the 713nt amlicon (c.1171+5G>A) shown in Figure 2D were gel extracted and Sanger sequenced. The WT product demonstrates canonical splicing between exons 7 and exon 8 while the c.1171+5G>A weakens the Exon 7 canonical donor splice site to generate the *GFAP-$\kappa$* isoform.
